# CircDNA-PAINT: high-resolution binary-barcode multiplexing with circular DNA docking strands

**DOI:** 10.64898/2026.11.14.766712

**Authors:** Junbo Wu, Xuejuan Chen, Longxiang Li, Yan Shi, Haijiao Xu, Hongda Wang, Yangang Pan

## Abstract

Understanding key biological processes governed by cell surface protein distribution requires highly multiplexed imaging. Binary barcode methods based on DNA hybridization represent a powerful approach to such imaging; however, higher multiplexing demands longer DNA barcodes, which substantially compromise localization resolution. Here we introduce circular DNA-based multiplexed DNA-PAINT (CircDNA-PAINT), a method that achieves a resolution better than 20 nm and thus enables robust binary-encoded multiplexing. This approach avoids the linear time overhead of adaptor-cycling DNA-PAINT and greatly shortens acquisition time for large-plex imaging. CircDNA-PAINT provides a new tool for dissecting complex nanoscale cellular architectures.

---

The spatial organization and interactions of biomolecules underpin biological function^1^. Spatial omics techniques hold great promise for this goal. With the aid of AI-driven image analysis, imaging- and mass spectrometry-based proteomics now enable the quantification of proteins at single-cell resolution ^2^, and DNA sequencing reads can be computationally assembled into spatial proteomics networks that currently resolve 76 proteins at the subcellular level^3^. However, many biomolecular interactions occur within tens of nanometers, far beyond the resolution of single-cell proteomics. Super-resolution microscopy offers a route to high resolution combined with multiplexed imaging. For example, secondary-label-based SUM-PAINT achieves virtually unlimited multiplexing at sub-15-nm resolution^4^; however, it requires toehold-mediated strand displacement after each imaging round and identifies only one protein type per round, imposing considerable time constraints. In contrast, binary barcode methods address 2^N^ − 1 species in N rounds, dramatically cutting the number of imaging cycles^5^. Nevertheless, a key limitation remains: achieving higher multiplexing requires longer DNA barcodes, which can substantially compromise localization resolution (**Fig. 1a**).

**Fig 1.**
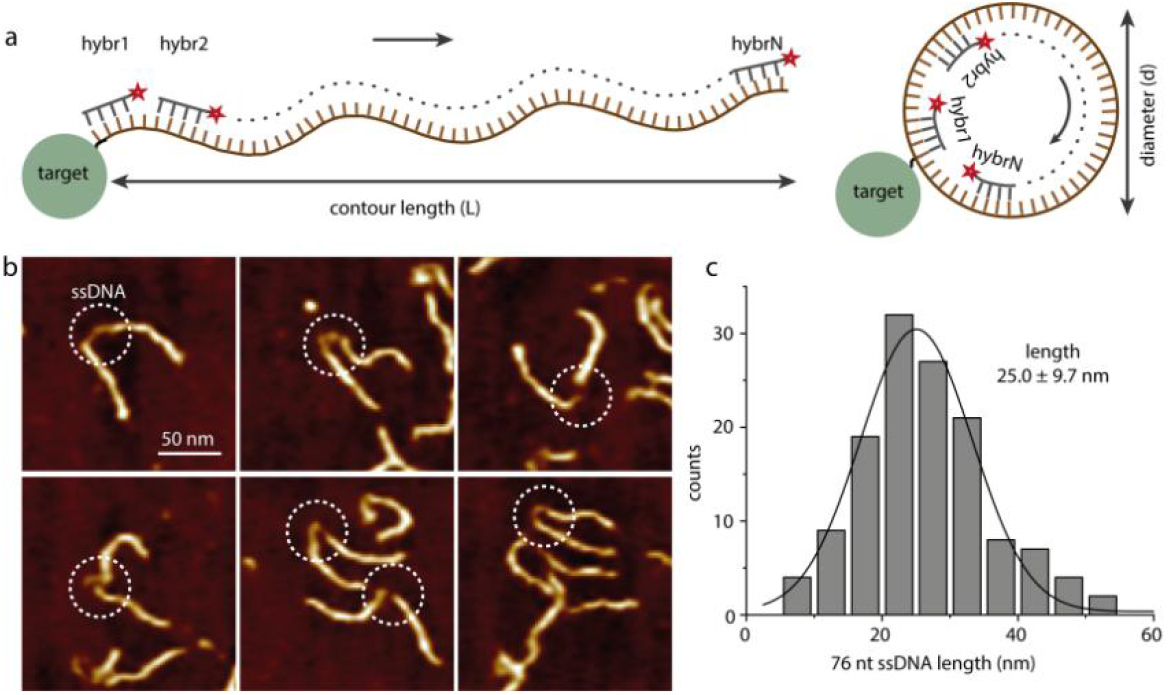
The concept of circular DNA-based PAINT. (a) Schematic of N rounds imaging with linear versus circular DNA. (b) AFM image of the 76 nt single-stranded DNA (highlighted by the white dashed circle). (c) Length distribution of the 76 nt ssDNA.

To surmount this challenge, we introduce a new concept: circular DNA-based multiplexed DNA-PAINT (CircDNA-PAINT). A linear docking strand of contour length L allows its free end to reach up to L from the anchor, spanning a region of diameter 2L; closing the same strand into a circle confines it to a loop of diameter L/π. This reduces the spatial extent from 2L to L/π, theoretically improving localization resolution by a factor of 2π ≈ 6, an idealized geometric upper bound (**Fig. 1a**). In addition, to reduce identification error, we adapted the MHD4 code previously developed for MERFISH^5^. In this scheme, each barcode has exactly four “1” bits and thus only four motifs available for imager binding (**Table S1**). To maintain imaging speed, we used four distinct 19-nt motifs^6^ (M1-M4; 76 nt in total), each containing multiple overlapping binding sites. Gap-DNA measurements showed that the fourth motif lies ∼25 nm from the anchor (**Fig. 1b, c, Supplementary Fig. 1**). Consistent with this, simulations gave a 95% containment radius of ∼22.7 nm for the far segment (**Supplementary Fig. 2a**). Together, these results suggest that, in principle, features separated by less than ∼40 nm cannot be resolved, because the far segment of the linear docking strand can reach ∼25 nm from its anchor and its localization cloud therefore spans roughly twice this distance. Rather than probing this boundary with a ∼40-nm grid, we used 20-nm grids, which were also used for the circular probes below. The linear docking strands indeed failed to resolve the 20-nm features (**Fig. 2a, b, Supplementary Fig. 3**). Notably, for circular DNA probes, the second and third motifs, rather than the fourth, have the lowest resolution; we therefore tested resolution using motif 2 instead of motif 4 (**Fig. 2c**). Interestingly, although the far segment of the 76-nt circle spans 38 nt, the same length as a 38-nt linear strand, its closed topology keeps the far-segment cloud more compact (**Supplementary Fig. 2**). Accordingly, the patterns could be clearly resolved (**Fig. 2d**). To test whether circularization affects hybridization kinetics, we also imaged motif 2 in its linear form. Quantitative PAINT (qPAINT) analysis yielded comparable binding events (249 versus 263) for the linear and circular structures (**Fig. 2e, f**). Together, these results demonstrate that circular DNA docking strands enable multiplexed imaging with a resolution better than 20 nm. Given that SUM-PAINT attains ∼6.6 nm average localization precision despite its long secondary labels^4^, circDNA-PAINT should likewise achieve such label-size-limited resolution.

**Fig 2.**
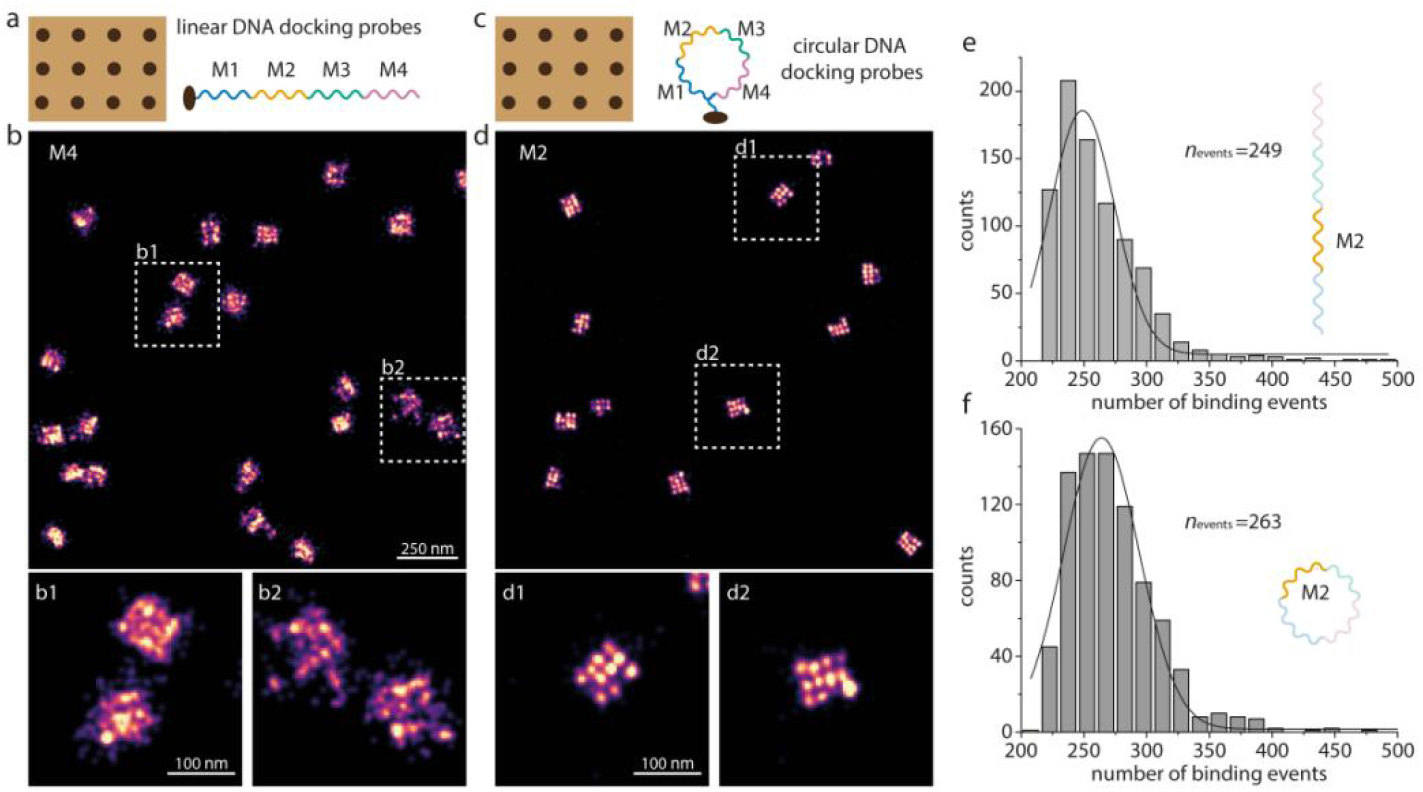
Comparison between the linear and circular docking probes. (a) Design of 20-nm origami grids carrying an extended linear single-stranded DNA (ssDNA). (b) DNA-PAINT images of the 20-nm origami grids obtained with an imager strand targeting motif 4 (M4); scale bar, 250 nm. (b1, b2) Enlarged views of the boxed regions in (b); scale bar, 100 nm. (c) Design of 20-nm origami grids carrying an extended circular ssDNA. (d) DNA-PAINT images of the 20-nm origami grids obtained with an imager strand targeting motif 2 (M2) of the circular probes; (d1, d2) Enlarged views of the boxed regions in (d); scale bar, 100 nm. (e, f) Distributions of binding events on entire DNA origami structures, obtained with an imager strand targeting M2, for (e) linear and (f) circular docking probes.

Two primary multiplexing strategies, SUM-PAINT^4^ and FLASH-PAINT^7^ originating from exchange-PAINT, have been established for super-resolution imaging. They perform one imaging cycle per target, consequently, imaging time scales nearly linearly with target count.

Binary-barcode DNA-PAINT offers an attractive alternative, where N rounds can theoretically encode 2^N^-1 distinct molecular species. Yet practical implementation has been hindered: longer linear docking barcodes expand the spatial reach of imager strands and degrade localization precision. CircDNA-PAINT overcomes this bottleneck using circular DNA docking strands. Combined with MHD4 error-correcting codes, only 12 imaging rounds are needed for 60 targets.

Because we used the same docking motifs as SUM-PAINT, the two schemes could be compared directly. SUM-PAINT relies on repeated cycles of secondary-adaptor hybridization (15 min), imaging acquisition (17 min), and toehold-mediated extinction/wash (15 min), requiring ∼60 h for 60-plex measurements. For binary barcoding, by contrast, we practically need to consider only the imaging time (17 min), so 60 targets can in theory be read out in ∼3 h. This breaks the linear time-multiplexing trade-off of sequential-cycling workflows.

We note that this work provides proof of principle on DNA origami. Further work will be required to adapt CircDNA-PAINT for complex cellular samples. Even so, our results show that circularized docking strands preserve localization precision at the barcode lengths required for high multiplicity. We further note that, in principle, our design could also increase imaging speed by adding concatenated repeats without sacrificing resolution, because a circular docking strand remains more compact than a linear strand of the same length. Together, these properties make CircDNA-PAINT an attractive molecular tool for high-multiplex nanoscale spatial proteomics.

## Materials

All the DNA oligonucleotides were purchased from Sangon Biotech. M13mp18 scaffold was obtained from New England Biolabs. BSA-biotin was ordered from Shenzhen Braveds Biotechnology Co., Ltd. (Shenzhen, China). Tween-20 was ordered from Biosharp (Hefei, China). Streptavidin was purchased from Thermo Fisher Scientific (Cat. No. 434302). PBS buffer was purchased from Biosharp (Hefei, China). EDTA was purchased from Gen-View / GENVIEW (Houston, TX, USA). TE buffer was obtained from Sangon Biotech (Shanghai) Co., Ltd. (Shanghai, China). Coverslips (22×50 mm and 22×22 mm) were purchased from Fisherbrand. Ninety-nanometer gold nanoparticles (G-90-100) were ordered from Cytodiagnostics.

The following buffers were used for sample preparation and imaging:

DNA origami assembly buffer: 1×TE buffer, 10 mM MgCl_2_

DNA origami purification buffer: 1×TE buffer, 10 mM MgCl_2_

DNA origami sample preparation buffer: 10 mM Tris pH 8, 100 mM NaCl and 0.05% Tween-20

DNA origami imaging buffer: 1×PBS, 1 mM EDTA and 500 mM NaCl, pH 7.4;

### DNA origami self-assembly

DNA origami self-assembly was carried out in a one-pot reaction system with a total volume of 100 μL^8^. The mixture contained 10 nM scaffold strand, 100 nM folding staples, 500 nM biotinylated staples, and 1 μM staple strands carrying docking-site extensions, all prepared in assembly buffer. The reaction solution was first ramped to 90 °C and held for 15 min, then cooled to 70 °C at a rate of 1 °C min^−1^, followed by a further decrease to 30 °C at 3 °C min^−1^, and finally maintained at 4 °C.

### DNA origami purification

DNA origami structures were purified using a 100 kDa ultrafiltration concentrator (Merck KGaA, Darmstadt, Germany). Briefly, 400 μL of deionized water was added to the concentrator, which was then centrifuged at 5,000 × g and 4 °C for 5 min. Next, 100 μL of DNA origami solution and 300 μL of DNA purification buffer were added, and the concentrator was centrifuged at 2,000 × g and 4 °C for 15 min; this centrifugation step was repeated twice. The filter was then inverted into a fresh tube and centrifuged at 1,000 × g and 4 °C for 2 min, and the recovered solution was collected and stored at −20 °C.

### DNA origami sample preparation and imaging

For sample preparation, a coverslip (22×50 mm) and a parafilm strip were bonded with UV-curable glue to form a chamber with an internal volume of approximately 50 µL. The chamber was first filled with 50 µL of biotin-labeled BSA (1.0 mg mL^−1^ in DNA origami sample preparation buffer) and incubated for 5 min, then rinsed with 200 µL of the same buffer. Subsequently, 50 µL of streptavidin (1.0 mg mL^−1^ in the same buffer) was introduced and allowed to bind for 5 min. After sequential washing with 200 µL each of the preparation buffer and the DNA origami purification buffer, 50 µL of biotin-labeled DNA structures (∼200 pM in purification buffer) was injected and incubated for another 5 min. The chamber was then washed with 200 µL of DNA origami imaging buffer, and finally, 100 µL of the imager solution (in imaging buffer) was flowed in. The chamber was subsequently sealed with a cover slide (22×22 mm).

Fluorescence imaging was performed on an inverted microscope (Nikon Instruments) equipped with an objective-type TIRF configuration and an oil-immersion objective (NA 1.49). Excitation was provided by a 642-nm diode-pumped solid-state (DPSS) laser (MPB Communications, 2 W). Images were acquired with an exposure time of 100 ms, an EM gain of 100, and an illumination power density of approximately 0.3 kW cm^−2^; a total of 40,000 frames were collected for reconstruction.

### Image analysis

Raw fluorescence data were subjected to super-resolution reconstruction using the Picasso software package^6^. Drift correction was performed with an adaptive intersection maximization (AIM). For quantification of binding kinetics, localizations were linked into trajectories with a maximum gap of three frames and a maximum distance of 130 nm^6^. The kinetic information of the selected structures was exported via the “save pick properties” command. Subsequent analyses, including histogram plotting and curve fitting, were performed in OriginPro.

## Data availability

All raw data are available upon reasonable request from the authors.

## Acknowledgements

This work was supported by National Key Research and Development Program of China (No. 2024YFA1307400), National Natural Science Foundation of China (22327808 to Hongda Wang, 22577129 to Yangang Pan), ‘100 Talents Project’ of Chinese Academy of Sciences, China (to Yangang Pan).

## Author contributions

Y.P., H.W., and X.C. conceived and designed the experiments. Y.P., J.W., and X.C. carried out the experimental work, performed data analysis, and contributed to manuscript preparation. Y.P. and H.W. supervised the overall study and data interpretation. Y.P. and X.C. wrote the paper. All authors edited the manuscript.

