## Supplemental file for "CircDNA-PAINT: high-resolution binary-barcode multiplexing with circular DNA docking strands"

**Supplementary Table 1 | An example of 31 molecular species using the MHD4 code.**

| Molecular Species | MHD4 code word | Signal rounds |
|---|---|---|
| molecular01 | 10101010000 | 1,3,5,7 |
| molecular02 | 00011100001 | 4,5,6,11 |
| molecular03 | 10000100011 | 1,6,10,11 |
| molecular04 | 00011001010 | 4,5,8,10 |
| molecular05 | 00110000011 | 3,4,10,11 |
| molecular06 | 10010000110 | 1,4,9,10 |
| molecular07 | 00000111010 | 6,7,8,10 |
| molecular08 | 10000101100 | 1,6,8,9 |
| molecular09 | 01011010000 | 2,4,5,7 |
| molecular10 | 00101100010 | 3,5,6,10 |
| molecular11 | 00110101000 | 3,4,6,8 |
| molecular12 | 01100001100 | 2,3,8,9 |
| molecular13 | 01010001001 | 2,4,8,11 |
| molecular14 | 01000010011 | 2,7,10,11 |
| molecular15 | 00000001111 | 8,9,10,11 |
| molecular16 | 00100100101 | 3,6,9,11 |
| molecular17 | 00001011100 | 5,7,8,9 |
| molecular18 | 11100000001 | 1,2,3,11 |
| molecular19 | 00111000100 | 3,4,5,9 |
| molecular20 | 11001000010 | 1,2,5,10 |
| molecular21 | 00100011001 | 3,7,8,11 |
| molecular22 | 01100110000 | 2,3,6,7 |
| molecular23 | 10001001001 | 1,5,8,11 |
| molecular24 | 10100001010 | 1,3,8,10 |
| molecular25 | 01001000101 | 2,5,9,11 |
| molecular26 | 00010110100 | 4,6,7,9 |
| molecular27 | 11010100000 | 1,2,4,6 |
| molecular28 | 11000010100 | 1,2,7,9 |
| molecular29 | 10010010001 | 1,4,7,11 |
| molecular30 | 01000100110 | 2,6,9,10 |
| molecular31 | 01001101000 | 2,5,6,8 |

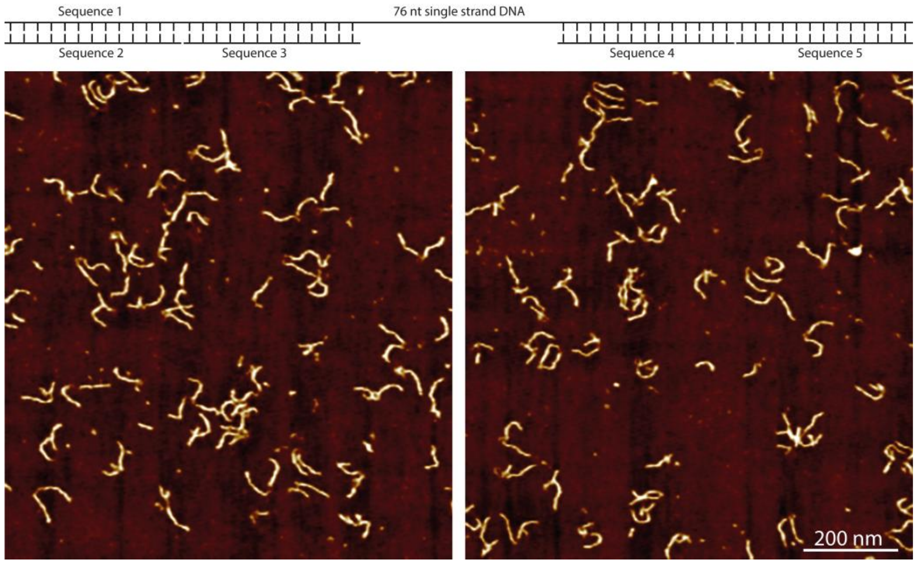


**Supplementary Figure 1 | Design and AFM image of DNA containing a 76-nt single-stranded region.**

**Supplementary Table 2 | Gap DNA sequence.**

| Name | Sequence |
|---|---|
| Sequence1 | AATGCTACTACTATTAGTAGAATTGATGCCACCTTTTCAGCTCGCGCCCCA AATGAAAATATAGCTAAACAGGTTATTGACCATTTGCGAAATGTATCTAAT GGTCAAACTAAATCTACTCGTTCGCAGAATTGGGAATCAACTGTTACATG GAATGAAACTTCCAGACACCGTACTTTAGTTGCATATTTAAAACATGTTG AGCTACAGCACCAGATTCAGCAATTAAGCTCTAAGCCATCCGCAAAAATG ACCTCTTATCAAAAGGAGCAATTAAAGGTACTCTCTAATCCTGACCTGTT GGAGTTTGCTTCCGGTCTGGTTCGCTTTGAAGCTCGAATTAAAACGCGAT ATTTGAAGTCTTTCGGGCTTCCTCTTAATCTTTTTGATGCAATCCGCTTTG CTTCTGACTATAATAGTCAGGGTAAAGACCTGATTTTTGATTTATGGTCATT CTCGTTTTCTGAACTGTTTAAAGCATTTGAGGGGG |
| Sequence2 | TGGTCAATAACCTGTTTAGCTATATTTTCATTTGGGGCGCGAGCTGAAAAG GTGGCATCA ATTCTACTAATAGTAGTAGCATT |
| Sequence3 | CAACATGTTTTAAATATGCAACTAAAGTACGGTGTCTGGAAGTTTCATTCC ATGTAACAGTTGATTCCCAATTCTGCGAACGAGTAGATTTAGTTTGACCAT TAGATACATTTCGCAAA |
| Sequence4 | CGGATTGCATCAAAAAGATTAAGAGGAAGCCCGAAAGACTTCAAATATC GCGTTTTAATTCGAGCTTCAAAGCGAACCAGACCGGAAGCAAACTCCAA CAGGTCAGGATTAGAGAGTACC |
| Sequence5 | CCCCCTCAAATGCTTTAAACAGTTCAGAAAACGAGAATGACCATAAATCA AAAATCAGGTCTTTACCCTGACTATTATAGTCAGAAGCAAAG |

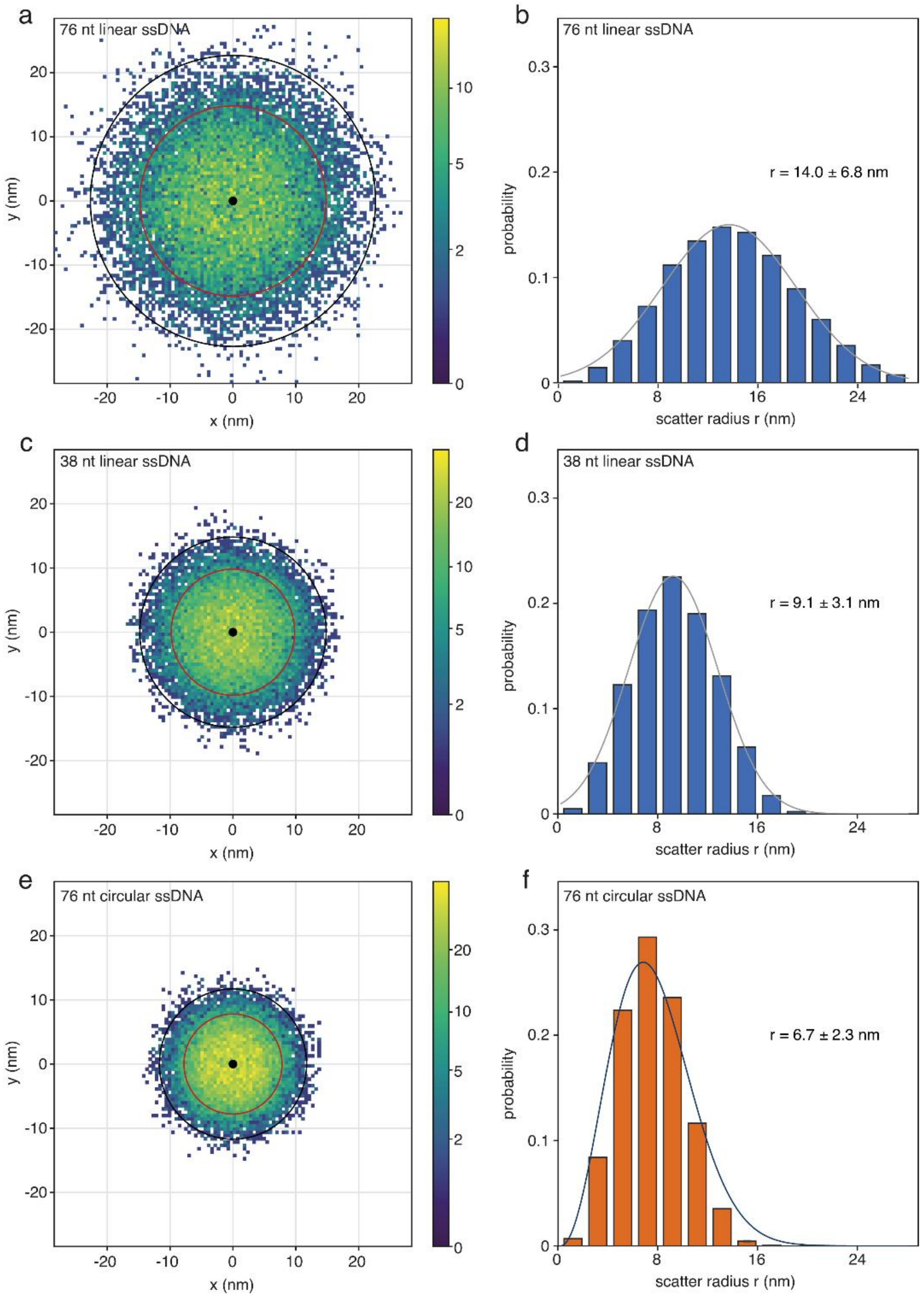


**Supplementary Figure 2 | Coarse-grained Monte Carlo simulations of the accessible regions of the far segment of tethered linear and circular single-stranded DNA (ssDNA), based on 32 independent Monte Carlo replicas.** (a) XY projection of the position cloud sampled by the far segment of a 76-nt linear ssDNA. The black circle indicates the 95% containment radius (22.7 nm), and the red circle indicates the root-mean-square distance from the anchored nucleotide. (b) Radial probability distribution of the free end, calculated in successive 2-nm shells; the distribution is broad, extends beyond 26 nm and peaks at approximately 14 nm. (c) XY projection of the position cloud sampled by the far segment of a 38-nt linear ssDNA. The black circle indicates the 95% containment radius (14.8 nm), and the red circle indicates the root-mean-square distance from the anchored nucleotide. (d) Radial probability distribution of the free end, calculated in successive 2-nm shells; the distribution peaks at approximately 9 nm. (e) XY projection of the position cloud sampled by the far segment of a 76-nt circular ssDNA. The black circle indicates the 95% containment radius (11.7 nm), and the red circle indicates the root-mean-square distance from the anchored nucleotide. (f) Radial probability distribution of the tracked far-segment site, calculated in successive 2-nm shells; the distribution peaks at approximately 7 nm. In all XY projections, colour encodes the number of positions per bin on a logarithmic scale, and all panels share the same axis ranges, enabling direct comparison of the three position clouds and their corresponding radial distributions.

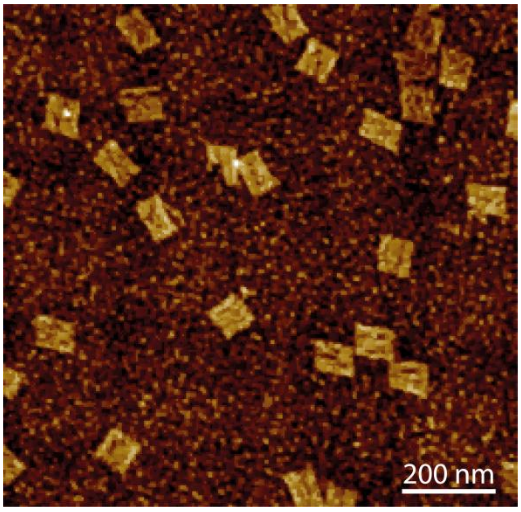


**Supplementary Figure 3** | AFM image of the synthesized DNA origami; Scale bar: 200 nm.

**Supplementary Table 3 | DNA-PAINT docking strand for experiments on a 20-nm grid.**

| | |
|---|---|
| Docking strand | TCCTCCTCCTCCTCCTCCTACCACCACCACCACCACCACTCTCTCTCT CTCTCTCTCACACACACACACACACACA |

**Supplementary Table 4 | Imager strand sequence.**

| Name | Sequence | 3'-modification |
|---|---|---|
| Imager2 | TGGTGGT | ATTO-647N |
| Imager4 | TGTGTGT | ATTO-647N |

**Supplementary Table 5 | Imaging parameters.**

| Dataset | Parameters |
|---|---|
| Figure 2b | 100 ms, 40000 frames, 500 pM Imager4, 300 W/cm² (642 nm) |
| Figure 2d | 100 ms, 40000 frames, 500 pM Imager2, 300 W/cm² (642 nm) |

### Supplementary Note

#### 1. Simulation system construction

We simulated three single-stranded DNA (ssDNA) constructs using the same coarse-grained model: a 76-nucleotide (nt) linear strand, a 38-nt linear strand, and a covalently closed 76-nt circular strand. The 76-nt linear strand comprised beads 0–75, with its 5′-terminal nucleotide (bead 0) serving as the anchor. The 38-nt linear strand comprised beads 0–37 and was constructed in the same manner. The circular strand was formed by connecting consecutive beads i and i + 1 for i = 0–74, together with a closing bond between beads 75 and 0. One nucleotide of the ring (bead 0) was arbitrarily selected as the anchor. All three systems used the same interaction parameters and categories of Monte Carlo moves. The free end of the 38-nt linear strand and the antipodal nucleotide of the 76-nt ring (bead 38) were separated from their respective anchors by 37 and 38 bonds, respectively.

#### 2. Coarse-grained model and force field

Each nucleotide was represented by a spherical bead. The chain was described by a discrete worm-like-chain Hamiltonian incorporating excluded-volume interactions:

$$U = U_{bond} + U_{bend} + U_{WCA}.$$

Adjacent beads interacted through a harmonic bond potential:

$$U_{bond} = (1/2)\ k_b\ (r - b_0)^2,$$

where r is the instantaneous bond length, $b_0 = 0.7$ nm is the effective contour-length increment per nucleotide for ssDNA [1,2], and $k_b = 100$ kT nm$^{-2}$. In the harmonic approximation, this stiffness corresponds to a bond-length standard deviation of $(kT/k_b)^{1/2} = 0.1$ nm.
Chain stiffness was represented by a bending potential acting on the angle θ between consecutive bond vectors:

$$U_{bend} = (1/2)\ \kappa\theta^2.$$

The bending constant κ was determined without using the small-angle approximation. Instead, it was obtained by numerically solving

$$\langle \cos\theta \rangle = \exp(-b_0/l_p),$$

the worm-like-chain tangent-correlation relation [3]. The angular average was evaluated using the full angular measure sin(θ) dθ over $0 \leq \theta \leq \pi$ rather than a Gaussian approximation. For $b_0 = 0.7$ nm and a persistence length $l_p = 1.5$ nm, this procedure yielded $\kappa = 1.949$ kT rad$^{-2}$. The persistence length $l_p = 1.5$ nm was selected as a representative value within the reported salt-dependent range for ssDNA [4,5]. Thus, the simulations represent an effective implicit-salt condition rather than a specific buffer composition.
Nonbonded beads separated by at least two nucleotides along the contour interacted through the Weeks–Chandler–Andersen (WCA) potential [6]:

$$U_{WCA}(r) = 4\varepsilon\ [(\sigma/r)^{12} - (\sigma/r)^6] + \varepsilon, \qquad r < 2^{1/6}\sigma,$$

and $U_{WCA}(r) = 0$ otherwise. The bead diameter was σ = 0.7 nm, and the interaction strength was ε = 1 kT. For the circular strand, contour separation was defined by the shorter of the two paths around the ring. Directly bonded pairs were excluded from the WCA interaction. No periodic spatial boundary conditions were applied.

For numerical stability in the independent Brownian-dynamics calculations, the WCA potential below $r_{cap}$ = 0.70 nm was replaced by its tangent line at $r_{cap}$. This linearization limits the pair force to approximately 34 kT $nm^{-1}$ while preserving the potential and force at $r_{cap}$. In a control Monte Carlo simulation, replacing the original WCA core with the linearized form changed all reported structural observables by less than 0.6%.

Water and ions were treated implicitly. The model did not explicitly include Coulomb interactions, hydrodynamic interactions, base pairing, base stacking, or sequence-dependent secondary structure. It therefore provides a polymer-level description of accessible conformations rather than a sequence-specific structural model. Coarse-grained DNA models that explicitly resolve base pairing and stacking are available when sequence-dependent secondary structure is essential[7].

### 3. Anchoring and coordinate system

Bead 0 was constrained to the origin throughout each simulation. After each accepted Monte Carlo move, the chain was translated rigidly to return bead 0 to the origin. This translation does not change the potential energy and is equivalent to fixing one nucleotide with an infinitely stiff, zero-length tether. All reported distances from the anchor were measured relative to this nucleotide. For the linear strands, the anchored nucleotide was the 5′ terminus; for the circular strand, the anchoring site was arbitrary because of the symmetry of the model.

### 4. Metropolis Monte Carlo sampling with mixed moves

Equilibrium conformations were sampled using a Metropolis Monte Carlo algorithm with three types of trial moves [8]. One sweep consisted of N attempted moves, where N is the number of nucleotides. At each attempt, a move type was selected with the following probabilities:

**Local displacement (0.55):** One nucleotide was displaced by a three-dimensional Gaussian vector. Each Cartesian component had a standard deviation of 0.20 nm, corresponding to a three-dimensional root-mean-square displacement of $\sqrt{3} \times 0.20 = 0.346$ nm.

**Crankshaft rotation (0.15):** One nucleotide was rotated by an angle uniformly sampled from $[0, 2\pi)$ about the axis defined by its two neighbors. This move preserves the lengths of both adjacent bonds[9,10].

**Pivot move (0.30):** For a linear chain, the terminal fragment beyond a randomly selected bead was rotated by a uniformly sampled angle about a random axis passing through that bead[11]. For the ring, a contiguous internal arc was rotated about the chord connecting its two bounding beads. The arc contained at least two and at most $N - 2$ bonds, ensuring that neither the moved arc nor its complement was trivial.

Each proposal scheme was symmetric, such that the forward and reverse moves had equal proposal probabilities. Trial moves were therefore accepted with the standard Metropolis probability $\min[1, \exp(-\Delta U/kT)]$, with the internal energy unit set to kT = 1. Only bonds, bending angles, and nonbonded pairs affected by a move were reevaluated when calculating $\Delta U$. For every tested move, the locally calculated energy change was checked against the difference in total energy. Because these Monte Carlo sweeps sample equilibrium conformations rather than physical dynamics, no physical time step or dynamical trajectory was inferred from them.

### 5. Equilibration and production protocol

Initial conformations were self-avoiding random coils constructed with approximately fixed bond lengths and a minimum separation of 0.75 $b_0$ between nonadjacent beads. Each replica underwent 3,000 equilibration sweeps followed by 15,000 production sweeps. One conformation was stored every 25 production sweeps, yielding 600 stored conformations per replica. Each system was simulated with 32 independent replicas using different random seeds, giving 19,200 stored conformations per system. All stored production conformations were included in the ensemble averages; no frames were discarded after equilibration.

The mean acceptance rates per attempted move were 0.383, 0.390, and 0.338 for the 76-nt linear, 38-nt linear, and 76-nt circular strands, respectively. To assess sensitivity to the initial conformation, additional control runs were initiated from a fully extended rod. The mean-square end-to-end distance differed by approximately 1.7% for both $\varepsilon = 0$ and $\varepsilon = 1$ kT, suggesting that the production ensembles were insensitive to initial conformation within the statistical precision of these controls.

Monte Carlo was used for production sampling because only equilibrium distributions were required. The longest chain relaxation mode evolves on a much longer timescale than the stable Brownian-dynamics integration step imposed by the stiff bond and excluded-volume potentials[12]. Brownian dynamics was therefore used only as an independent cross-check, as described below.

### 6. Independent Brownian-dynamics cross-check

An independent overdamped Brownian-dynamics implementation used the same model parameters, with the short-range WCA linearization described in Section 2, and the Euler–Maruyama scheme[13]:

$$x_i(t + dt) = x_i(t) + dt\, F_i(t) + (2dt)^{1/2}\, \xi_i(t),$$

where $F_i$ is the force on bead i and $\xi_i$ is a three-dimensional Gaussian random vector with zero mean and unit variance in each Cartesian component. The equation is expressed in reduced units: lengths are measured in nm, energies in kT, and time in $\tau_0 = 1\ \mathrm{nm}^2/D_0$. The model nucleotide diffusion coefficient was set to $D_0 = 0.3\ \mathrm{nm}^2\ \mathrm{ns}^{-1}$, giving $\tau_0 = 3.33$ ns. The anchored bead was returned to the origin after each integration step. For each topology, 24 independent trajectories were initiated from equilibrated Monte Carlo conformations selected from different parts of the stored trajectories. Each trajectory comprised $2 \times 10^6$ steps with $dt = 5 \times 10^{-4}\ \tau_0 = 1.67$ ps, corresponding to 3.33 μs. Configurations were stored every 1,000 steps, yielding 2,000 frames per trajectory.

Time-step sensitivity was assessed by repeating 12 linear-chain trajectories over the same physical interval using $dt = 2.5 \times 10^{-4}\ \tau_0$. Halving the time step changed the mean-square end-to-end distance from 201.6 to 203.8 $\mathrm{nm}^2$, a difference of 1.1%. The Brownian-dynamics and Monte Carlo ensembles agreed within 0.5% for the bond-vector correlation and within 0.5% for the radius of gyration of the circular chain. The free-end distance from the anchor for the linear chain was 11% smaller in the finite-length Brownian-dynamics trajectories, consistent with incomplete sampling of the slowest relaxation mode. Accordingly, Brownian dynamics was used to cross-check the force field and local equilibrium statistics, whereas all reported accessible-region statistics were calculated from the Monte Carlo ensemble.

### 7. Definition of observables and statistical analysis

The principal observable was the position r of the distal segment relative to the anchored nucleotide. For the 76-nt linear strand, the distal segment was the free 3′ end (bead 75, 75 bonds from the anchor). For the 38-nt linear strand, it was the free 3′ end (bead 37, 37 bonds from the anchor). For the 76-nt circular strand, it was the antipodal nucleotide (bead 38, 38 bonds from the anchor along either arc). Thus, the circular distal segment and the free end of the 38-nt linear strand had nearly matching contour separations of 38 and 37 bonds, respectively.

For each system, we calculated the root-mean-square distance $R_{rms} = \langle r^2 \rangle^{1/2}$, the 50th (median), 90th, 95th, and 99th percentile distances $r_q$, the three principal standard deviations of the positional distribution, the radius of gyration $R_g$, the maximum distance of any nucleotide from the anchor, and, for the ring, the maximum pairwise distance (diameter). The 95% containment volume was defined as

$$V_{95} = (4/3)\pi r_{95}^3,$$

and the equivalent local concentration was calculated as

$$c_{eff} = 1/(N_A V_{95}),$$

where $N_A$ is Avogadro's constant and $V_{95}$ is converted from $nm^3$ to liters. This quantity represents an occupancy-based equivalent concentration for the distal segment within a spherical containment region; it should not be interpreted as a kinetic association rate. Accessible-volume and effective-concentration concepts have been applied to tethered nucleic acids and FRET-restrained structural models [14,15]. We also calculated the root-mean-square distance from the anchor as a function of contour separation s, $R_s = \langle |r(s)|^2 \rangle^{1/2}$, for every nucleotide along each chain.

Unless otherwise stated, central values were first averaged within each independent replica and then combined across the 32 replicas. Uncertainties are standard errors of the mean across replicas. Percentile and containment-volume estimates were calculated from pooled production conformations, with replica-level uncertainties reported where applicable. Radial distributions were summarized by empirical histograms and least-squares fits to an ideal Gaussian-chain radial form and an empirical smooth Gaussian. These fits were descriptive only and did not affect the simulation parameters.

### 8. Reproducibility and model scope

All simulations and analyses were performed using a purpose-written Python/NumPy implementation. Monte Carlo replicas used random seeds 1000 + r, and Brownian-dynamics replicas used seeds 5000 + r, where r denotes the replica index. The stored conformations, analysis tables, scripts, and random-seed conventions are provided with this work. Because base pairing and base stacking were not explicitly modeled, the predicted accessible regions should be interpreted as those of an unstructured polymer rather than as sequence-specific conformational distributions. Salt effects were represented only through the selected persistence length, and no explicit topological knot constraint was imposed on the circular strand.


1 Chi, Q., Wang, G. & Jiang, J. The persistence length and length per base of single-stranded DNA obtained from fluorescence correlation spectroscopy measurements using mean field theory. *Physica A: Statistical Mechanics and its Applications* **392**, 1072-1079, doi:https://doi.org/10.1016/j.physa.2012.09.022 (2013).

2 Tinland, B., Pluen, A., Sturm, J. & Weill, G. Persistence Length of Single-Stranded DNA. *Macromolecules* **30**, 5763-5765, doi:10.1021/ma970381+ (1997).

3 Fredrickson, G. H. The theory of polymer dynamics. *Current Opinion in Solid State and Materials Science* **1**, 812-816, doi:https://doi.org/10.1016/S1359-0286(96)80106-9 (1996).

4 Murphy, M. C., Rasnik, I., Cheng, W., Lohman, T. M. & Ha, T. Probing Single-Stranded DNA Conformational Flexibility Using Fluorescence Spectroscopy. *Biophysical Journal* **86**, 2530-2537, doi:https://doi.org/10.1016/S0006-3495(04)74308-8 (2004).

5 Chen, H. *et al.* Ionic strength-dependent persistence lengths of single-stranded RNA and DNA. **109**, 799-804, doi:10.1073/pnas.1119057109 (2012).

6 Weeks, J. D., Chandler, D. & Andersen, H. C. Role of Repulsive Forces in Determining the Equilibrium Structure of Simple Liquids. *The Journal of Chemical Physics* **54**, 5237-5247, doi:10.1063/1.1674820 (1971).

7 Ouldridge, T. E., Louis, A. A. & Doye, J. P. K. Structural, mechanical, and thermodynamic properties of a coarse-grained DNA model. *The Journal of Chemical Physics* **134**, 085101, doi:10.1063/1.3552946 (2011).

8 Metropolis, N., Rosenbluth, A. W., Rosenbluth, M. N., Teller, A. H. & Teller, E. Equation of State Calculations by Fast Computing Machines. *The Journal of Chemical Physics* **21**, 1087-1092, doi:10.1063/1.1699114 (1953).

9 Verdier, P. H. & Stockmayer, W. H. Monte Carlo Calculations on the Dynamics of Polymers in Dilute Solution. *The Journal of Chemical Physics* **36**, 227-235, doi:10.1063/1.1732301 (1962).

10 *Understanding Molecular Simulation: From Algorithms to Applications*. (Academic Press, Inc., 1996).

11 Madras, N. & Sokal, A. D. The pivot algorithm: A highly efficient Monte Carlo method for the self-avoiding walk. *Journal of Statistical Physics* **50**, 109-186, doi:10.1007/BF01022990 (1988).

12 Rouse, P. E., Jr. A Theory of the Linear Viscoelastic Properties of Dilute Solutions of Coiling Polymers. *The Journal of Chemical Physics* **21**, 1272-1280, doi:10.1063/1.1699180 (1953).

13 Ermak, D. L. & McCammon, J. A. Brownian dynamics with hydrodynamic interactions. *The Journal of Chemical Physics* **69**, 1352-1360, doi:10.1063/1.436761 (1978).

14 Woźniak, A. K., Schröder, G. F., Grubmüller, H., Seidel, C. A. M. & Oesterhelt, F. Single-molecule FRET measures bends and kinks in DNA. **105**, 18337-18342, doi:10.1073/pnas.0800977105 (2008).

15 Kalinin, S. *et al.* A toolkit and benchmark study for FRET-restrained high-precision structural modeling. *Nature methods* **9**, 1218-1225, doi:10.1038/nmeth.2222 (2012).
